# Di-Ras2 Promotes Renal Cell Carcinoma Formation by Activating the MAPK Pathway in the Absence of VHL

**DOI:** 10.1101/683821

**Authors:** Hanyu Rao, Xuefeng Li, Min Liu, Jing Liu, Xiaoxue Li, Jin Xu, Li Li, Wei-Qiang Gao

## Abstract

Clear cell renal cell carcinoma (ccRCC) is one of the most common and lethal human urological malignancies in the world. The pathological drivers for ccRCC are still poorly understood. One of them is the Ras family of small GTPases that function as “molecular switches” in many diseases including ccRCC. Among them, Di-Ras2 encodes a 26-kDa GTPase that shares 60% homology to Ras and Rap. Yet little is known about the biological function(s) of Di-Ras2. In this study, we found that Di-Ras2 was upregulated in ccRCC, and promoted the proliferation, migration and invasion of human ccRCC cells in the absence of von Hippel-Lindau (VHL). Mechanistically, Di-Ras2 induces and regulates ccRCC formation by modulating phosphorylation of the downstream effectors and activating the MAPK signaling pathway. Moreover, Di-Ras2 interacts with E3 ubiquitin ligase, VHL, which facilitates the ubiquitination and degradation of Di-Ras2. Together, these results indicate a potential function of Di-Ras2 as an oncogene and biomarker in ccRCC, and these data provide a new perspective of the relationship between VHL and the MAPK pathway in ccRCC tumorigenesis.

## Introduction

Renal cell carcinoma (RCC) originates from the renal tubular epithelium and is the most common type of kidney cancer, accounts for >90% of cancers in the kidney [1]. RCC is also the 7^th^ an 9^th^ most common cancer in men and women, respectively, worldwide, and accounts for approximately 2–3% of all adult malignancies [2–4]. Major subtypes with ≥5% incidence are clear cell RCC (ccRCC) [5], papillary RCC (pRCC) [6] and chromophobe RCC (chRCC) [7]. The most common subtype of RCC is clear cell renal cell carcinoma (ccRCC), which accounts for 75–80% of all diagnosed cases [8]. Early diagnosis of the disease is difficult, and metastases often occur before the primary tumor is detected. Therefore, radical nephrectomy is considered to be the most effective way to treat ccRCC [9, 10]. But the five-year survival rate for individuals with metastatic disease is still less than 10% [11]. It has been reported that hepatocyte growth factor (HGF) binding to its receptor, tyrosine-protein kinase Met, leads to activation of the Ras/MAPK signaling pathway to promote ccRCC growth and metastasis [9, 12]. Function of the RAS-Mitogen activated protein kinase (MAPK) signaling pathway (also known as the RAS-RAF-MEK-ERK pathway) is to integrate extracellular signals and to coordinate a suitable response by a subsequent control of cellular growth, survival, and differentiation[13]. Aberrant activation of this pathway is a major and highly prevalent oncogenic event in many human cancers including ccRCC. Therefore, how Ras/MAPK signaling pathway is activated is important for us to understand the molecular mechanism of ccRCC, and suppression of Ras/MAPK signaling cascade may be one of the strategies to treat ccRCC.

The function of MAPK kinase is mediated by Ras GTPase signaling in normal and disease states [14, 15]. As an important component in Ras/MAPK signaling pathway, the small GTPase Ras family regulates a wide range of cell functions, including proliferation, differentiation, migration and invasion[16, 17]. The GTP-binding Ras-like protein 2 (Di-Ras2) is one of GTPases forming a distinct subgroup of the Ras family, and shares 30–40% overall identity with other members of the Ras family [18, 19]. Di-Ras2 forms a tight complex with SmgGDS in cytosol, which lowers its affinity for guanine nucleotides [20]. Analyses using Oncomine and The Cancer Genome Atlas (TCGA) databases show that mRNA expression levels of Di-Ras2 are significantly up-regulated in some tumors, especially ccRCC. However, little is known about the biological role(s) of Di-Ras2 or how their activities are regulated in ccRCC.

ccRCC is genetically characterized by a frequent loss or mutation of von Hippel-Lindau (VHL) tumor suppressor gene, which appears to be the most important gene for the formation and progression of ccRCC [1, 5, 21].The tumor-suppressive effect of VHL protein comes from its function as an E3 ubiquitin ligase complex substrate recognition element [22, 23].Moreover, VHL takes part in the ubiquitin-proteasome pathway (UPP) to control steady-state protein level of several effectors in ccRCC[24, 25]. Although Hypoxia-inducible factor α-subunit (HIF) is the most studied gene interacting with VHL[26–28], recent studies also showed that VHL can interact with several other oncogenic proteins and facilitates their ubiquitination and degradation in ccRCC, such as Jade family PHD finger 1 (Jade-1)[29–31],Zinc fingers and homeoboxes 2 (ZHX2)[32], protein phosphatase 5 (PP5)[33], Aurora kinase A (AURKA)[34], and atypical protein kinase C (PKC)[35]. Moreover, VHL has also been reported to regulate the epigenetic landscape of renal cancer by binding and degrading the hyperphosphorylated Rpb1 [36]. Therefore, according to its multifunction, von Hippel-Lindau (VHL) is highly related to ccRCC by enhancing the ubiquitination and degradation of certain oncogenes. However, whether and how VHL can directly interact with the MAPK pathway is unclear.

In the present study, we sought to characterize the physiological properties of Di-Ras2 proteins in ccRCC. To this end, we employed a plasmids-mediated overexpression and knockdown system to determine Di-Ras2’s function in ccRCC. We found that Di-Ras2 promoted ccRCC proliferation by activating the MAPK pathway in the absence of von Hippel-Lindau (VHL), which can facilitate the ubiquitination and degradation of Di-Ras2. We thus propose that Di-Ras2 is a potential oncogene and biomarker in ccRCC, and these data provide us a new view of the relationship of VHL and the MAPK pathway in ccRCC tumorigenesis.

## Results

### Di-Ras2 expression is up-regulated in ccRCC

To determine the significance of Di-Ras2 in ccRCC, we first analyzed multiple microarray data sets in the Oncomine and TCGA databases. There were about 100 genes differentially expressed in human kidney tumor samples as compared to the adjacent normal kidney tissues. Among them, Di-Ras2 exhibited the most significant up-regulation in ccRCC samples (Supplementary Fig. S1). As shown in Figs. 1A-F, Di-Ras2 mRNA levels were increased significantly in human tumor samples as compared to the adjacent normal kidney tissues.

**Figure 1.**
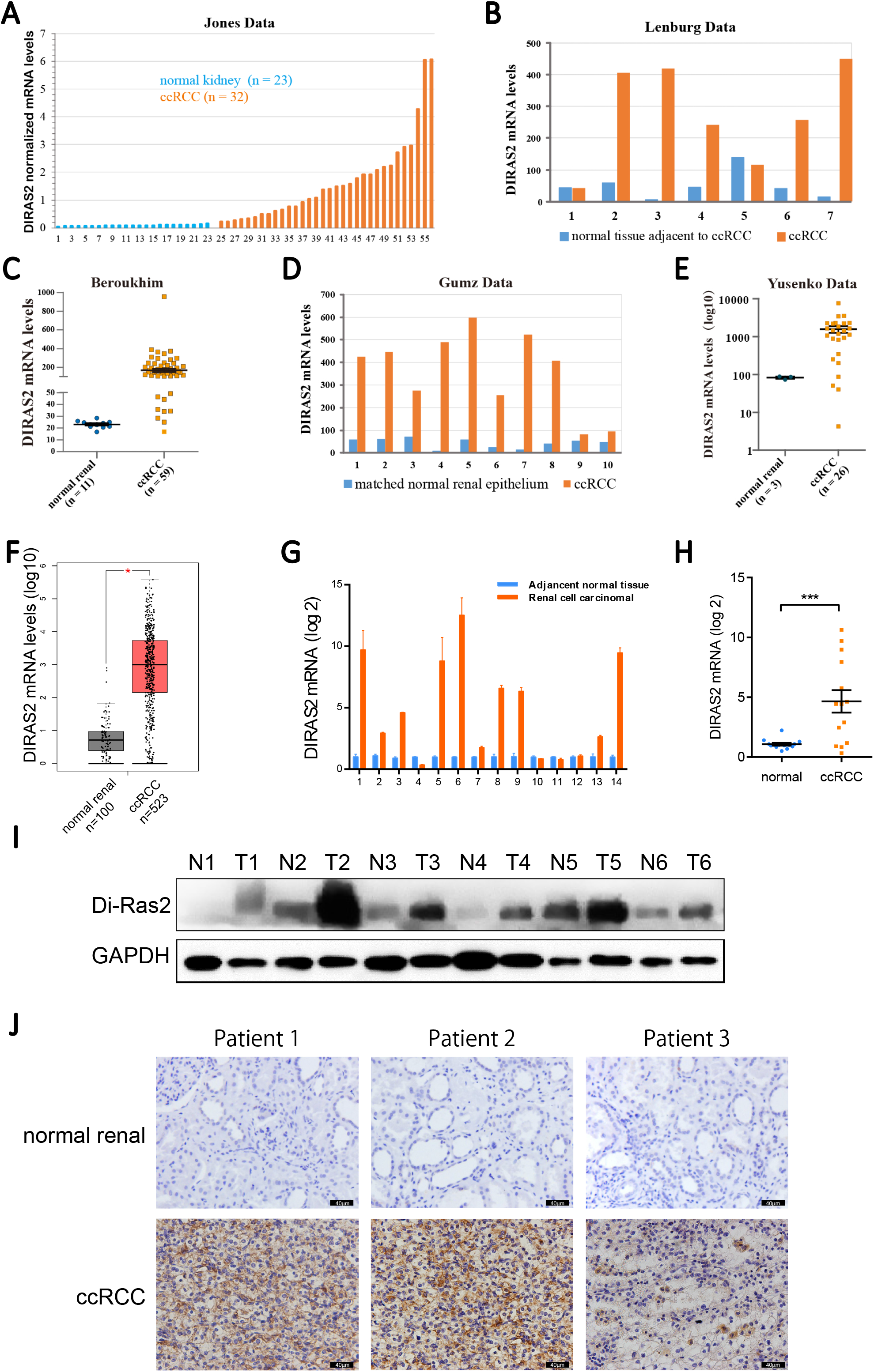
Di-Ras2 expression is up-regulated in ccRCC. (**A-F**) Analysis from the Oncomine and TCGA databases showed that mRNA expression levels of Di-Ras2 were significantly higher in renal cell carcinoma compared with normal tissues. (**G-H**) Di-Ras2 mRNA expression level in 14 paired tumor samples and normal tissue. (**I**) Immunoblotting showed higher protein levels of Di-Ras2 in 5 of 6 tumor samples compared with the respective matched normal tissues (T, tumor; N, normal tissue). (**J**) Immunohistochemical staining on normal and renal cell carcinoma tissues with anti-Di-Ras2 antibody.

To verify the microarray analysis results, we performed q-PCR, Western blotting, and IHC staining on human ccRCC specimens and their matched normal tissues. Data showed that mRNA and protein expression levels of Di-Ras2 were significantly higher in renal cell carcinoma compared with normal tissues in most of the paired samples. (Figs. 1G-J).

### Di-Ras2 promotes VHL-mutated ccRCC tumor growth *in vitro* and *in vivo*

To examine the biological function of Di-Ras2 in ccRCC, we then employed lentivirus-mediated overexpression and knockdown systems in 786-O cells (Figs. 2A and 2B), which has been established as one of the first RCC cell lines with many characteristics of ccRCC, and is used most commonly in RCC-focused research [37]. As shown in Figs. 2C and 2D, cell proliferation was increased significantly by overexpression of Di-Ras2 and was suppressed by knockdown of Di-Ras2 in 786-O cells. In addition, results from cell wound scratch assays and matrigel invasion trans-well assays demonstrated that Di-Ras2 overexpression promoted cell migration and invasion abilities, and Di-Ras2 knockdown attenuated the migration and invasion abilities of 786-O cells (Figs. 2E and 2F). Cell cycle analysis (Figs. 2G and 2H) revealed the same phenotype. Next, we employed lentivirus-mediated overexpression and knockdown systems in CAKI-1 cells (Figs. S2A and S2B), which is a widespread model line of ccRCC harboring wild-type VHL [37]. Surprisingly, there was no significant change on its proliferation (Figs. S2C and D), migration and invasion abilities (Supplementary Figs. S2E and S2F) after overexpression and knockdown of Di-Ras2 in CAKI-1 cells. Cell cycle analysis also revealed that Di-Ras2 had no effect on of CAKI-1 cells (Supplementary Figs. S2G and S2H). Moreover, we examined another VHL-mutated model of ccRCC, A498, and another VHL-wild-type RCC cell line, ACHN [37]. Results from A498 cells and ACHN cells showed similar phenotypes with 786-O cells and CAKI-1 cells respectively (Supplementary Figs. S3 and S4). These data suggest that Di-Ras2 promotes tumor progression in ccRCC cell lines in the absence of VHL.

**Figure 2.**
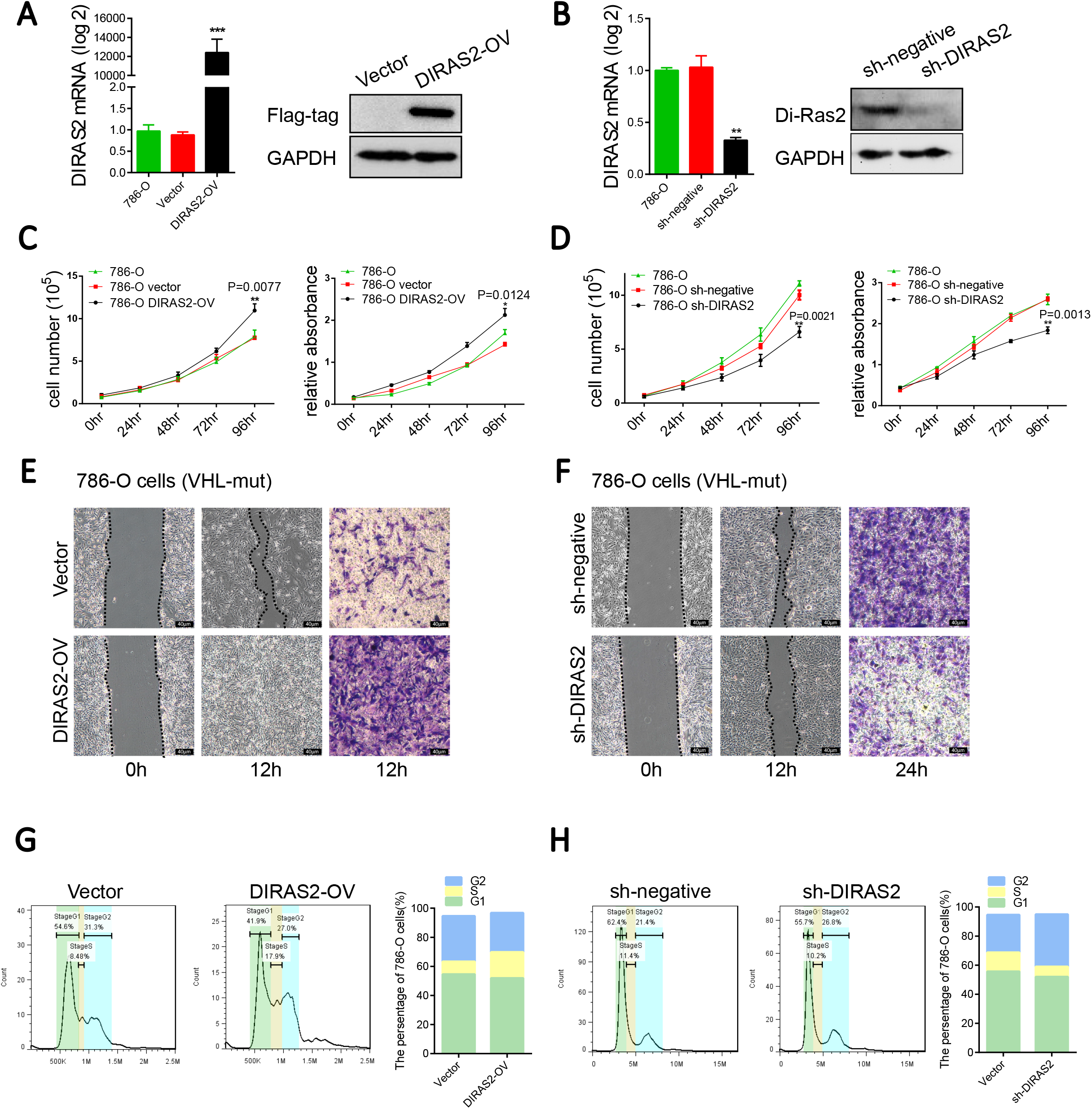
Di-Ras2 promotes ccRCC cell proliferation, migration, and invasion in VHL-mutated cell lines. mRNA and protein levels of Di-Ras2 after (**A**) overexpression and (**B**) knockdown of Di-Ras2 in 786-O cells lines. (**C, D**) Cell proliferation abilities were compared with their vector control cells. Cell migration and invasion abilities of (**E**) DIRAS2-OV 786-O cells as well as (**F**) DIRAS2-KD 786-O cells were compared with their vector control cells, as demonstrated by cell wound scratch and trans-well assays. The percentages of (**G**) DIRAS2-OV 786-O cells as well as (**H**) DIRAS2-KD 786-O cells in the G1, S, and G2 phases for each sample are shown. VT, vector; WT, wild-type; MUT, mutated; OV, overexpression; KD, knockdown. Data are shown as the mean value ± SEM.

**Figure 3.**
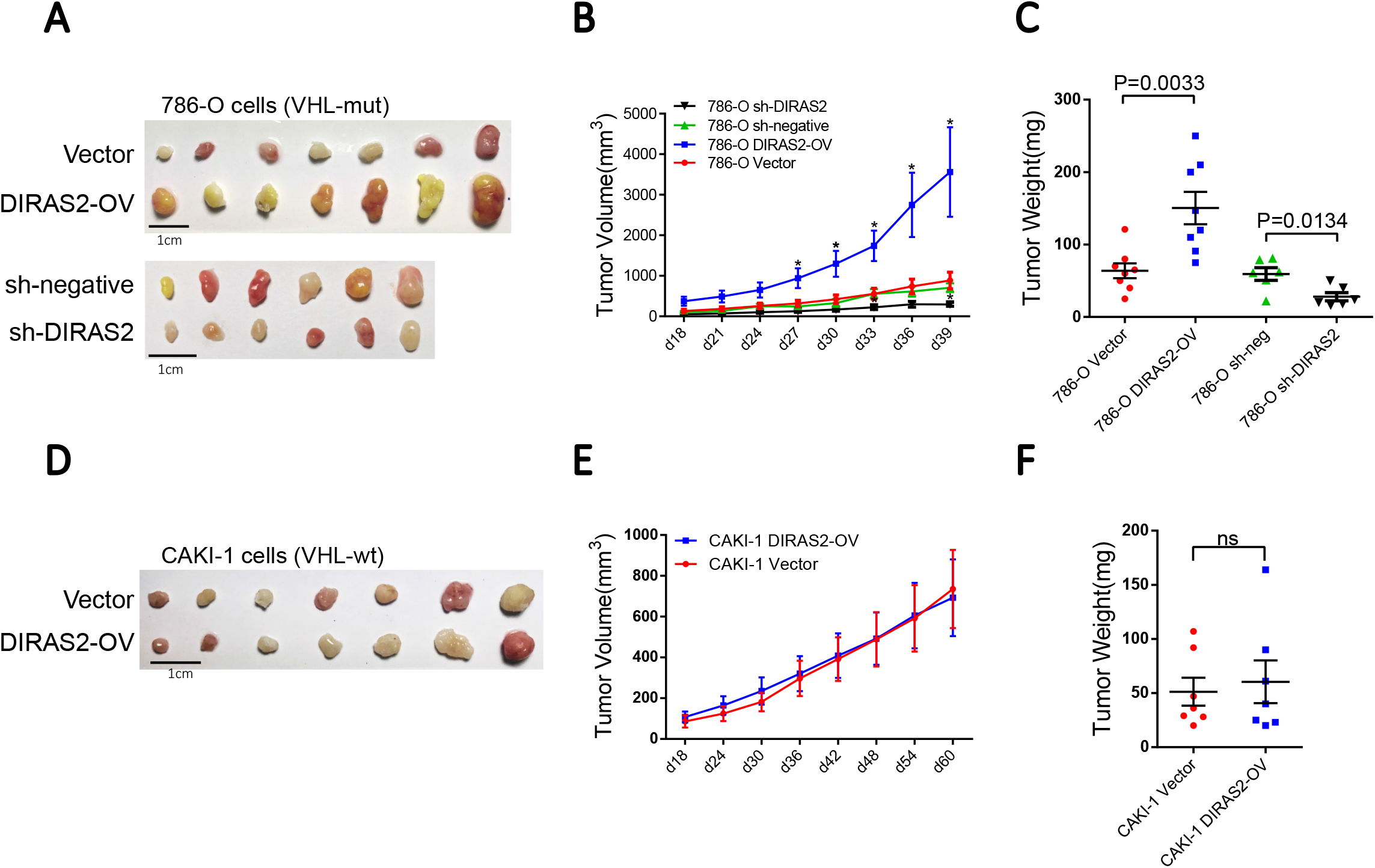
Di-Ras2 promotes VHL-mutated ccRCC tumor growth *in vivo*. (**A**) Xenograft tumor assays using 786-O cells stably transfected by DIRAS2-overexpression and sh-DIRAS2 lentiviruses. (**B**) Tumor volumes of xenografts generated by DIRAS2-OV and DIRAS2-KD 786-O cells were compared with their vector control cells. (**C**) Tumor weights of xenografts generated by DIRAS2-OV and DIRAS2-KD 786-O cells were compared with their vector control cells. (**D**) Xenograft tumor assays using CAKI-1 cells stably transfected by DIRAS2-overexpression lentiviruses. (**E**) Tumor volumes of xenografts generated by DIRAS2-OV CAKI-1 cells were compared with their vector control cells. (**F**) Tumor weights of xenografts generated by DIRAS2-OV CAKI-1 cells were compared with their vector control cells. Data are shown as the mean value ± SEM.

**Figure 4.**
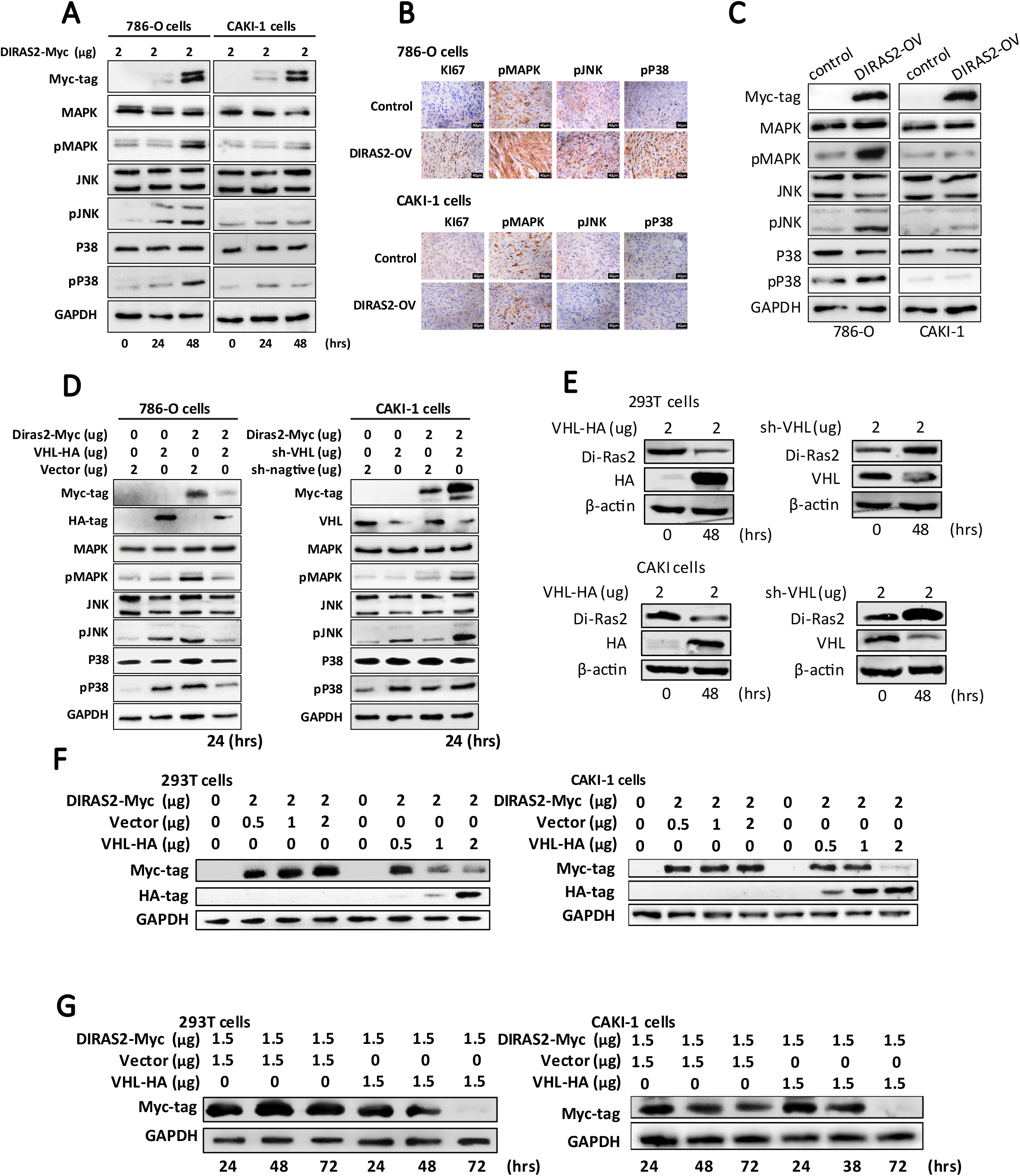
Di-Ras2 activates the MAPK pathway in the absence of VHL. (**A**) Overexpression of Di-Ras2 activated the MAPK pathway in VHL-mut 786-O cells, but not VHL-wt CAKI-1 cells. (**B**) IHC on xenograft tumor sections. (**C**) WB on xenograft tumor samples. (**D**) VHL attenuated the function of Di-Ras2 in activating MAPK in 786-O cells, and the deficiency of VHL restored the function of Di-Ras2 in activating MAPK in CAKI-1 cells. (**E**) The protein level of endogenous Di-Ras2 after overexpression and knockdown of VHL in 293T and CAKI-1 cells (48h). (**F**) The protein level of Di-Ras2 decreased in tandem with the progressively increasing levels of VHL in 293T and CAKI-1 cells (24h). (**G**) The expression of VHL reduced the protein abundance of co-transfected Di-Ras2 in 293T cells and CAKI-1 cells.

To verify the stimulative role of Di-Ras2 in ccRCC progression *in vivo*, we performed xenograft tumor assays using 786-O and CAKI-1 cells stably transfected by Di-Ras2-overexpression and Di-Ras2-knockdown lentiviruses. Lentiviral expression of Di-Ras2 resulted in an accelerated xenograft tumor growth and bigger tumor volumes, and lentiviral knock down of Di-Ras2 resulted in an attenuated xenograft tumor growth and smaller tumor volumes in 786-O cells (Figs. 3A-C). However, Di-Ras2 overexpression produced no significant change in CAKI-1 cells (Figs. 3D-F). These data collectively indicate that Di-Ras2 acted as a potential tumor-promoting factor that positively regulates VHL-mutated ccRCC tumor growth.

### Di-Ras2 accelerates ccRCC tumor growth by activating the MAPK pathway in the absence of VHL

Given the fact that the Ras/MAPK pathways are often epigenetically dysregulated in ccRCC and MAPK kinase cascade functions as an essential effector cascade required for Ras GTPase signaling, we hypothesized that Di-Ras2may promote tumor growth through the MAPK pathway. Indeed, in the absence of VHL, Di-Ras2 promoted the phosphorylation of several MAPK pathway signal factors, such as MAPK, JNK, and P38 in 786-O cells. However, the MAPK pathway could not be activated by Di-Ras2 in CAKI-1 cells (wild type VHL) (Fig. 4A). In addition, we performed IHC and WB on xenograft tumor sections and samples as indicated in Fig. 3, and obtained the same results (Figs. 4B and 4C). These data confirmed that Di-Ras2 can promote the cascade reaction in the MAPK pathway in VHL-mutated ccRCC. To further validate the suppressive role of VHL in the activation of the MAPK pathway induced by Di-Ras2, we overexpressed VHL in 786-O cells and knocked down VHL in CAKI-1 cells. As shown in Fig. 4D, Di-Ras2 activated the MAPK signaling pathway in VHL-mutated 786-O cells, while this function was blocked by exogenous VHL. Meanwhile, the activation of MAPK signaling pathway was restored after knockdown of VHL in Di-Ras2-transfected CAKI-1 cells (wild type VHL). Surprisingly, we found that protein levels of both exogenous and endogenous Di-Ras2 was decreased when levels of VHL were elevated and was increased when levels of VHL were declined (Figs. 4D and 4E).

To further substantiate the suppressive role of VHL in the regulation of Di-Ras2, we investigated ectopic VHL expression on Di-Ras2 in 293T cells and CAKI-1 cells. In line with our hypothesis, protein levels of Di-Ras2 reduced proportionally when VHL levels were progressively augmented, suggesting that VHL negatively regulates Di-Ras2 abundance in a dose-dependent and time-dependent manner (Figs. 4F and 4G).

### VHL interacts with Di-Ras2 and enhances the ubiquitination and degradation of Di-Ras2

Increasing evidence indicates that the E3 ligase plays an important role in oncogenesis and cancer development by controlling protein levels that regulate multiple biological processes [38]. Given that VHL is a type of E3 ligase, we next investigated the molecular mechanism through which VHL promoted Di-Ras2 degradation. As shown in Fig. 5A, the half-life of Di-Ras2 protein was shortened significantly in 293T and CAKI-1 cells transfected with VHL after using cycloheximide (CHX), indicating that VHL negatively affects Di-Ras2 stability. In addition, as shown in Fig. 5B, we observed that treatment with MG132, a proteasome inhibitor, led to a significant lengthening of the Di-Ras2 protein half-life, supporting the notion that VHL regulates Di-Ras2 stability by proteasome-mediated mechanisms.

**Figure 5.**
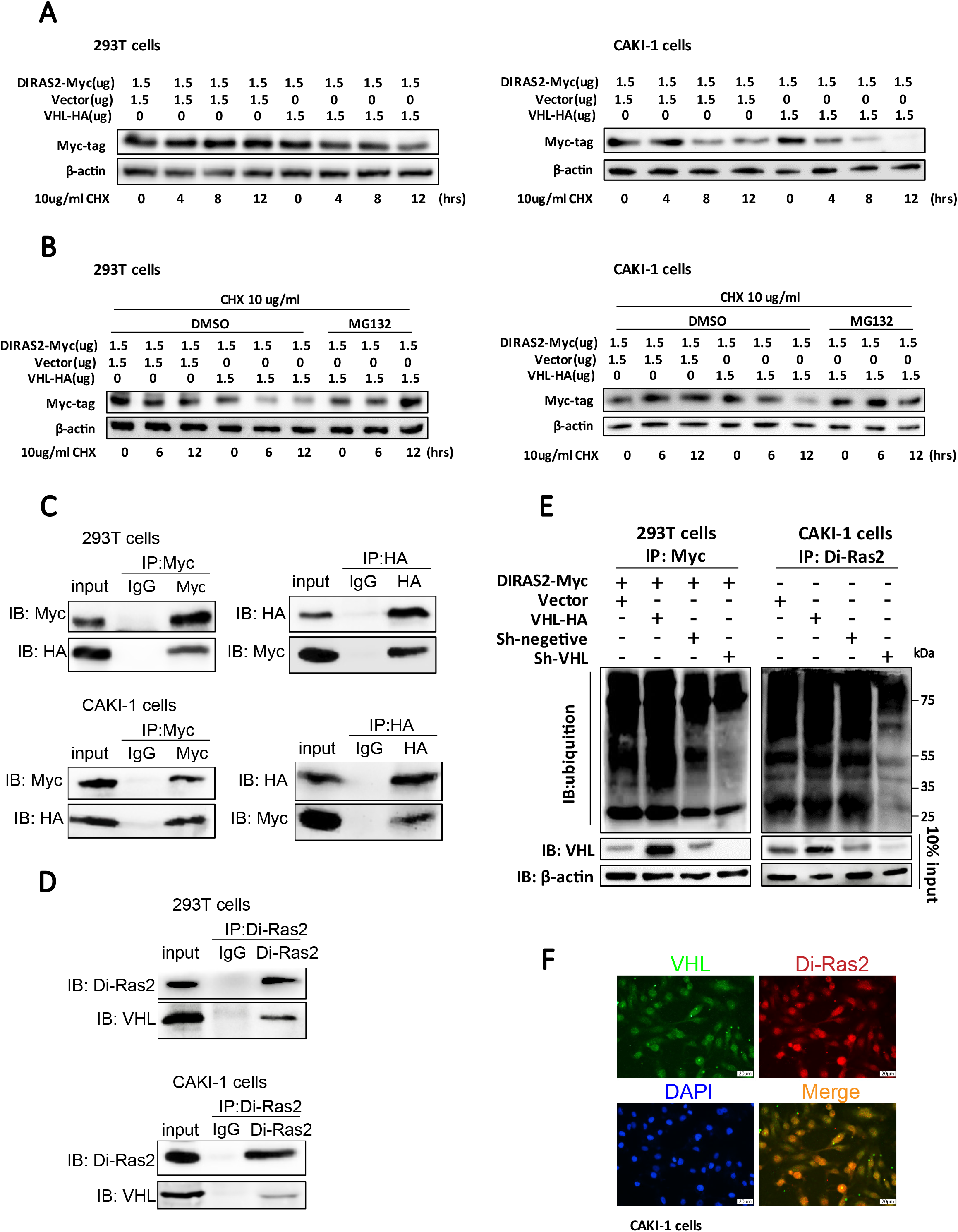
VHL binds to Di-Ras2 and facilitates the ubiquitination and degradation of Di-Ras2. (**A**) The half-life of Di-Ras2 was shortened by exogenous VHL in 293T cells and CAKI-1 cells. (**B**) The half-life of Di-Ras2 in 293T cells and CAKI-1 cells transfected with VHL was prolonged by MG132 treatment. (**C**) Di-Ras2 co-immunoprecipitated with VHL in 293T and CAKI-1 cells transfected with Myc-tagged Di-Ras2 and HA-tagged VHL. (**D**) Endogenous VHL or Di-Ras2 from protein complexes was detected. (**E**) Ubiquitination of exogenous and endogenous Di-Ras2 were enhanced by VHL overexpression and was attenuated by VHL knockdown in 293T cells and CAKI-1 cells. (**F**) Immunofluorescent staining on CAKI-1 cells with anti-Di-Ras2 and anti-VHL antibodies.

Given that protein-protein interaction is essential for E3 ligase to act on its substrates, we next explored whether VHL interacts with Di-Ras2. Co-immunoprecipitation results showed that both exogenous and endogenous VHL bound to Di-Ras2 in 293T cells and CAKI-1 cells (Figs. 5C and 5D). Considering that VHL protein serves as an E3 ligase, we hypothesized that VHL promoted Di-Ras2 protein ubiquitination. In agreement with our hypothesis, we observed that the ubiquitination of both exogenous and endogenous Di-Ras2 was augmented by VHL overexpression, whereas they were diminished by VHL knockdown in 293T cells and CAKI-1 cells respectively (Fig. 5E). To confirm the interaction between VHL and Di-Ras2, we employed immunofluorescent staining on Di-Ras2 and VHL. The results showed that they were co-localized in CAKI-1 cells. (Fig. 5F). Collectively, our data suggest that VHL binds to Di-Ras2 and enhances its ubiquitination and degradation.

### The tumor-promoting activity of Di-Ras2 is overridden by VHL

To verify the suppressive role of VHL in Di-Ras2-induced ccRCC progression, we then employed a lentivirus-mediated overexpression system of VHL on Di-Ras2 overexpressed 786-O (Di-Ras2-OV 786-O), and a lentivirus-mediated knockdown system of VHL on Di-Ras2 overexpressed 786-O (Di-Ras2-OV CAKI-1). Cell proliferation was suppressed by overexpression of VHL in Di-Ras2-OV 786-O cells (Fig. 6A). In addition, results from cell wound scratch assays and matrigel invasion trans-well assays demonstrated that VHL overexpression attenuated cell migration and invasion abilities which was increased by Di-Ras2-OV in 786-O cells (Fig. 6B). Meanwhile, cell proliferation was increased by knockdown of VHL in Di-Ras2-OV CAKI-1 cells (Fig. 6C). And VHL knockdown restored the migration and invasion abilities of Di-Ras2-OV CAKI-1 cells (Fig. 6D). We also performed xenograft tumor assays using these cell lines. Overexpression of VHL resulted in an attenuated xenograft tumor growth and smaller tumor volumes in Di-Ras2-OV 786-O cells (Figs. 6E and 6F), and knockdown of Di-Ras2 resulted in an accelerated xenograft tumor growth and bigger tumor volumes in Di-Ras2-OV CAKI-1 cells (Figs. 6G and 6H). These data suggest that the tumor-promoting activity of Di-Ras2 is overridden by VHL and Di-Ras2 promotes ccRCC tumor progression in the absence of VHL.

**Figure 6.**
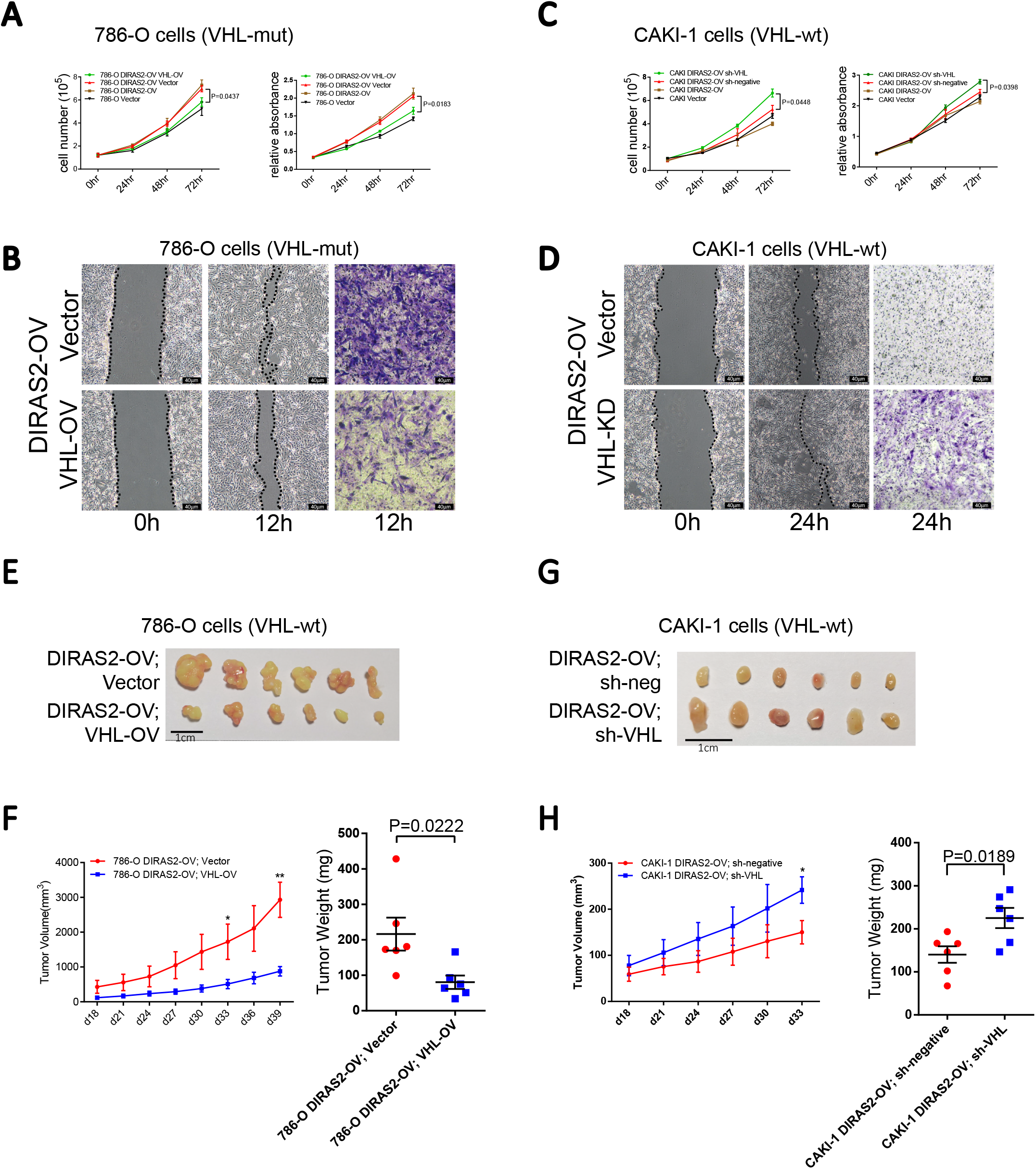
The tumor-promoting activity of Di-Ras2 is overridden by VHL. (**A**) Cell proliferation abilities of DIRAS2-OV 786-O cells after exogenous VHL transfection were compared with their vector control cells. (**B**) Cell migration and invasion abilities of DIRAS2-OV 786-O cells after exogenous VHL transfection were compared with their vector control cells, as demonstrated by cell wound scratch and trans-well assays. (**C**) Cell proliferation abilities of DIRAS2-OV CAKI-1 cells after sh-VHL transfection were compared with their vector control cells. (**D**) Cell migration and invasion abilities of DIRAS2-OV CAKI-1 cells after sh-VHL transfection were compared with their vector control cells, as demonstrated by cell wound scratch and tans-well assays. (**E**) Xenograft tumor assays using DIRAS2-OV 786-O cells stably transfected by VHL-overexpression lentiviruses. (**F**) Tumor volumes and weights of xenografts generated by VHL-OV; DIRAS2-OV 786-O cells were compared with their vector control cells. (**G**) Xenograft tumor assays using DIRAS2-OV CAKI-1 cells stably transfected by sh-VHL lentiviruses. (**H**) Tumor volumes and weights of xenografts generated by sh-VHL; DIRAS2-OV CAKI-1-1 cells were compared with their vector control cells. Data are shown as the mean value ± SEM.

We also tested the Di-Ras2 stability in hypoxia condition, which is closely related to VHL. The results showed that the ubiquitination of Di-Ras2 was augmented by VHL overexpression in 293T cells under both normal and hypoxia culture (Supplementary Figs. S5A and S5B).

### DiRas2 inversely correlates with VHL in ccRCC samples

Given our observations that Di-Ras2 can upregulate the MAPK pathway in human ccRCC and that VHL negatively regulated Di-Ras2 protein stability, we re-analyzed the TCGA RPPA data and found that the phosphorylation level of p38MAPK was significantly up-regulated in VHL-mut and DIRAS2-high ccRCC samples, which implicates an activation of the MAPK pathway in these samples (Fig. 7A). Further analyses showed a weak inverse correlation between the mRNA levels of Di-Ras2 and that of VHL in ccRCC samples from TCGA database (Fig. 7B). We also compared the Di-Ras2 and VHL protein levels from surgically removed human kidney tumor samples and RCC cell lines, using western blots and immunohistochemical staining, and found a clear inverse correlation between Di-Ras2 and VHL protein levels (Figs. 7C-E), which indicated that protein-protein interaction should play a dominant role in the downregulation of Di-Ras2. These findings suggested that the deficiency of VHL in ccRCC appear to promote the stability of Di-Ras2, leading to MAPK pathway activation, resulting in an accelerated tumor growth and progression (Fig. 7F).

**Figure 7.**
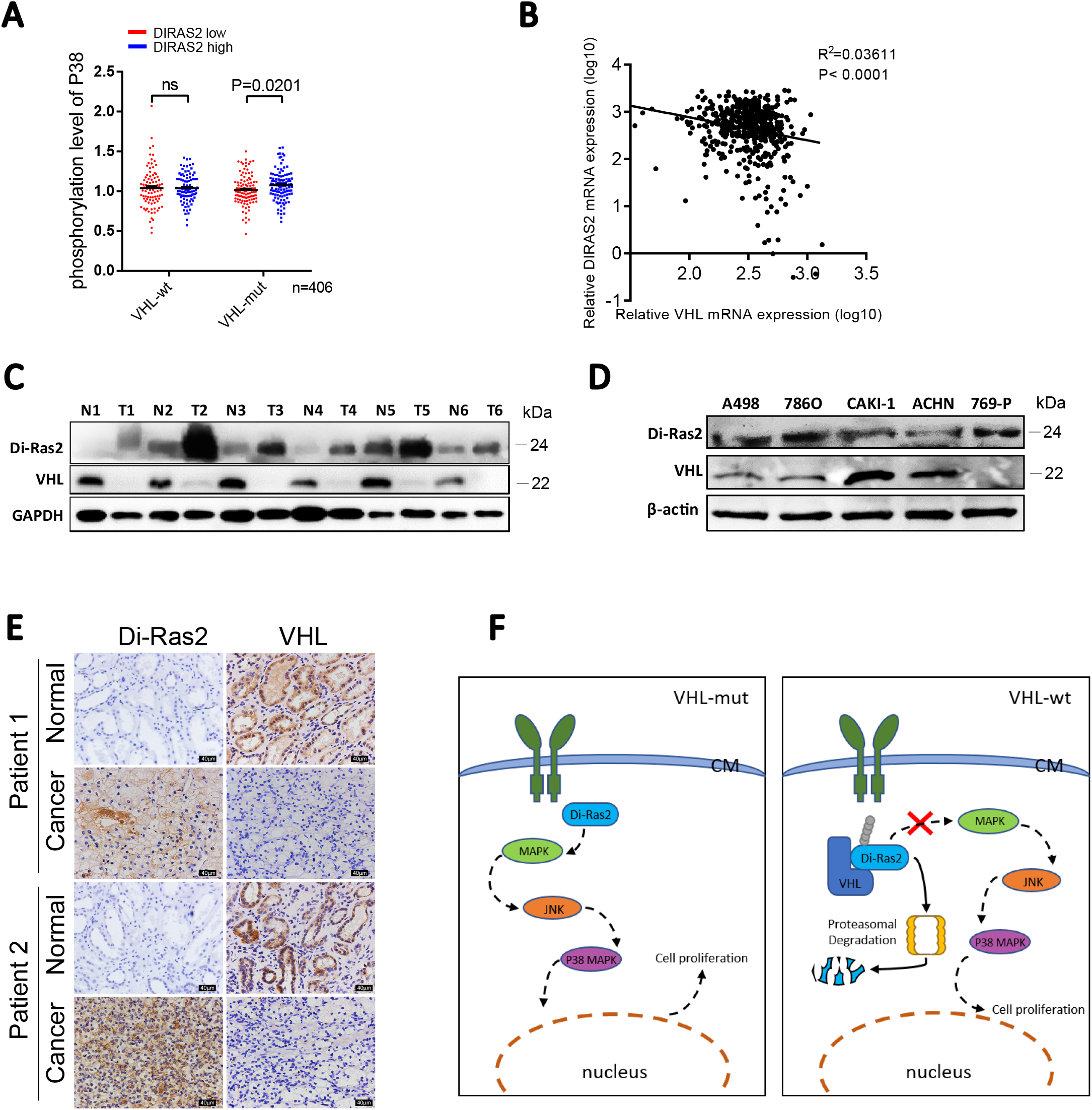
Di-Ras2 inversely correlates with of VHL in ccRCC samples. (**A**) Phosphorylation level of p38MAPK correlated with expression level of Di-Ras2 in VHL-mut or VHL-wt ccRCC samples (n=406). (**B**) Correlation between the expression levels of Di-Ras2 and VHL in TCGA ccRCC samples. (n=606). Western blotting of (**C**) ccRCC samples and (**D**) ccRCC cell lines with anti-Di-Ras2 and anti-VHL antibodies. (**E**) Immunohistochemical staining on normal and renal cell carcinoma tissues with anti-Di-Ras2 and anti-VHL antibodies. (**F**) Schematic representing the role of Di-Ras2 in tumorigenesis of ccRCC.

## Discussion

The small GTPase Ras family proteins are appreciated as potential candidates in many cancers. For example, the Ras/MAPK signaling pathway is altered in pancreatic ductal adenocarcinoma (PDAC), melanoma, lymphoblastic leukemia, lung adenocarcinomas, colon and rectal cancer including mutations in NRAS, KRAS or BRAF [39–44]. Moreover, the current study reinforces the idea that Ras/MAPK signaling pathway is activated predominantly with ccRCC [9, 12]. For patients with surgically resectable ccRCC, the standard of care is surgical excision by either partial or radical nephrectomy with a curative intent. By contrast, those with inoperable ccRCC typically undergo systemic treatment with targeted agents and/or immune checkpoint inhibitors [1, 10]. Accordingly, Ras/MAPK signaling pathway could be a promising target for ccRCC treatment. As a member of the Ras subfamily, yet little is known about the biological function(s) of Di-Ras2 in cancer research. Based on the MAPK signaling activation and its association with VHL, we have uncovered a specific function of Di-Ras2 in ccRCC as an activator of MAPK signaling pathway in the absence of VHL, which can facilitate the ubiquitination and degradation of Di-Ras2 and block its function in ccRCC. Our results thus implicate that Di-Ras2 is a novel oncogene which contributes to the formation of ccRCC and a potential target of VHL, linking the kidney-specific VHL tumor suppressor pathway and the MAPK signaling cascade in ccRCC.

VHL is a multifunctional protein and has been closely associated with ccRCC. To date, the most well-characterized function of VHL is its role as an E3 ubiquitin ligase that participates in the degradation of some oncogenic effectors. Of the several targets mentioned above, the regulation of the protein stability of HIFs is the most recognizable activity of VHL [35, 36, 45]. β-catenin can be down-regulated by VHL through stabilizing Jade-1 [29]. ZHX2 gene is often enriched in ccRCC cells in which VHL is lost or mutated [32, 46]. We showed in current study a potential target of VHL, Di-Ras2, and revealed a role of VHL in the regulation of ccRCC formation through the Ras/MAPK pathway. VHL physically binds to Di-Ras2 and enhances its ubiquitination and degradation. The loss of VHL results in dramatic changes in regulation of Di-Ras2, which can activate the MAPK pathway and induce ccRCC. This observation for the first time demonstrates that VHL can inhibit the MAPK signaling pathway through interaction with Di-Ras2.

Taken together, our findings have demonstrated the biological and clinical significance of Di-Ras2 as a potential oncogene in ccRCC. Di-Ras2 exerts an activating effect on the MAPK pathway and its downstream signals, thereby promoting ccRCC cells proliferation, migration and invasion. Meanwhile, its activity is regulated by VHL, which can promote the ubiquitination and degradation of Di-Ras2. In addition, our data suggest that activation of the MAPK pathway by Di-Ras2 represents a novel tumor promoter axis for VHL/Di-Ras2/MAPK. Furthermore, these findings directly link the kidney-specific VHL tumor suppressor pathway and the MAPK signaling cascade, and enlarge the scope of basic research on ccRCC formation. For clinical translation, pharmaceutical investigation of the interaction between Di-Ras2 and VHL may provide a potential promising strategy to inactivate MAPK signaling in ccRCC.

## Materials and Methods

### Patient samples

Microarray data from the Oncomine and TCGA databases were used. Twenty fresh samples of human ccRCC and paired normal tissue were obtained during surgery at the Department of General Surgery, Ren Ji Hospital. All samples were collected with the informed consent of patients and approved by the Animal Care committee of Shanghai Ren Ji Hospital. All cDNA and protein samples were extracted from those fresh tissue samples.

### RNA isolation and quantitative RT-PCR

Total RNA was isolated from cultured cells or fresh samples with Trizol reagent (Invitrogen). cDNA was synthesized by reverse transcription using the Prime Script RT reagent kit (TaKaRa) and subjected to quantitative RT-PCR with Di-Ras2, VHL, and ACTB or GAPDH primers in the presence of the SYBR Green Realtime PCR Master Mix (Thermo). Relative abundance of mRNA was calculated by normalization to ACTB or GAPDH mRNA. Data were analyzed from three independent experiments and were shown as the mean ± SEM.

### Immunoblotting

Cells were lysed with 100–300μL RIPA buffer supplemented with protease and phosphatase inhibitors (Millipore). The protein concentration was measured with the BCA Protein Assay (Bio-Rad). From each sample, 20–50 μg of total protein was separated by 8–12% SDS-PAGE gels and transferred onto nitrocellulose membranes (GM). Membranes were blocked in 5% BSA in TBS for 1 hour at room temperature, and then incubated with primary antibodies overnight at 4°C, washed in TBS containing 1% Tween20, incubated with an HRP-conjugated secondary antibody for 1 hour at room temperature, and developed by ECL reagent (Thermo). The immunoblots were quantified by Bio-Rad Quantity One version 4.1 software. Primary antibodies against Di-Ras2 (Proteintech ag7926), VHL (SAB #32075), and ubiquitin (sc-8017) were purchased from Proteintech, Signal way Antibody, and Santa Cruz Biotechnology. Antibodies against total-MAPK (#4695), p-MAPK (#4370), total-JNK (#9252), p-JNK (#4668), total-P38 (#9212), p-P38 (#4511), Flag (#14796), Myc-tag(#2276), HA-tag(#5017), GAPDH(#5174), and ACTB (#3700),were purchased from Cell Signaling Technology Inc.

### Immunofluorescent and immunohistochemistry (IHC) assays

To prepare slides of ccRCC cell lines for immunofluorescent staining, cells were plated on cover slides in 24-well plates and allowed to grow for 24 hours. The slides were then fixed in 4% formalin for 15 minutes at 4°C and rinsed by PBS three times. The cover slides were first treated with 0.5% Triton for 10 minutes and blocked with 10% goat serum at room temperature for 1 hour. Primary antibody incubations were performed overnight at 4°C. After extensive washing with PBS, secondary antibody was applied to the sections at room temperature for 1 hour. Slides were washed with PBS three times and then mounted with Vectashield mounting medium (Vector laboratories, Inc. H-1200). Primary antibody against Di-Ras2 (Abcam, ab67430) and VHL (SAB #32075) were used.

For IHC staining, paraffin-embedded ccRCC tissues were deparaffinized, rehydrated, and subjected to a heat-induced epitope retrieval step by treatment with 0.01M sodium citrate (pH 6.0). Endogenous peroxidase activity was blocked with 0.3% (v/v) hydrogen peroxide in distilled water. The sections were then incubated with 0.3% Triton X-100 in PBS (137 mM NaCl, 2.7 mM KCl, 10 mM Na_2_HPO_4_, 2 mM KH_2_PO_4_, pH 7.4) for 15 minutes, followed by 10% goat serum in PBS for 1 hour. Subsequently, samples were incubated with a mouse polyclonal Di-Ras2 antibody (Abcam, ab67430), diluted at 1:500 in 1% goat serum for 1 hour at 37°C. After three washes in PBS, sections were incubated with an HRP-conjugated secondary antibody for 1 hour at room temperature. Sections were counterstained with hematoxylin. Three random immunostaining images of each specimen were captured using a Leica DM2500 microscope and analyzed by Image-Pro Plus 6.0 software.

### Immunoprecipitation and ubiquitination assays

Cells were washed with ice-cold PBS and lysed in RIPA lysis buffer (50 mmol/L Tris-HCl, pH 8.0; 150 mmol/L NaCl; 1% NP-40) supplemented with protease and phosphatase inhibitors (Millipore) at 24 hours after transfection. Cell lysates were incubated with primary antibodies overnight at 4°C. Pierce™ Protein A/G Magnetic Beads (Thermo) were then added and the lysates were incubated for another 4 hours at 4°C. The immunoprecipitates were washed 4 times with the lysis buffer and boiled for 5 minutes at 98°C in protein loading buffer. Immunoprecipitated proteins were detected by subsequent immunoblotting. Antibodies used in the co-immunoprecipitation experiments were as follows: anti-Di-Ras2 (Proteintech ag7926), anti-VHL (SAB #32075), anti-Myc-tag(CST #2276), anti-HA(CST #5017), mouse immunoglobulin G, rabbit immunoglobulin G (Santa Cruz), glyceraldehyde-3-phosphate dehydrogenase (CST), and anti-ubiquitin (Santa Cruz sc-7199) antibodies.

### Protein half-life detection

HEK293T and CAKI-1 cells were co-transfected with the Di-Ras2-Myc expression plasmid and the VHL-Flag expression plasmid or the empty vector as described earlier. A total of 10 μg/mL of cycloheximide (CHX, purchased from Sigma) was added to the culture medium 24 hours after the plasmid transfection to 293T cells and CAKI-1 cells. Cells were lysed in RIPA buffer containing protease and phosphatase inhibitors as described earlier, after CHX treatment at indicated time points. For MG132 treatment, at indicated hours after transfection, cells were incubated with MG132 (10 μmol/L) for an additional 6 or 12 hours. Cells were then collected for immunoblots to determine the amount of Di-Ras2 protein.

### Cell lines and culture

All cell lines were obtained from the American Type Culture Collection (ATCC) and were confirmed by specific indices. CAKI-1, 786-O, A498, and 293T were cultured in RPMI-1640 or DMEM supplemented with 10% FBS (Thermo), 100U/mL penicillin, and 0.1 mg/mL streptomycin (Thermo) at 37°C in humidified 5% CO_2_ atmosphere. Hypoxic conditions were achieved with a hypoxia chamber (Billups-Rothenberg) flushed with a gas mixture of 1% O2, 5% CO2 and 94% N2.

### Plasmids, transfection, and lentivirus

ShRNA sequences for VHL and scramble shRNA were cloned into lentiviral vector pLVET-tTR-KRAB (GCCTGAGAATTACAGGAGACTCGAGTCTCCTGTAATTCTCAGGC). shRNA sequences for Di-Ras2 and scramble shRNA were cloned into lentiviral vector pLT3REVIR(AAGGTATATTGCTGTTGACAGTGAGCGACAGGATTGTTCTGTTCTAAAATAGTGAAGCCACAGATGTATTTTAGAACAGAACAATCCTGCTGCCTACTGCCTCGGACTTCAAGGGGCTA).

Human Di-Ras2 and VHL cDNA was generated by polymerase chain reaction and cloned into pCMV6-Entry vector with Myc-tag and HA-tag, respectively. Then, the cDNA of Di-Ras2 and VHL were cloned into the lentiviral expression vector pLenti.CMV.Puro.DEST.

All the constructs generated were confirmed by DNA sequencing. For transient transfection, cells were transfected with the jetPRIME® transfection reagent (Polyplus) according to the manufacturer’s instruction. Lentiviral packaging plasmids pCMV-DR8.8 and pMD2.G were co-transfected with the backbone plasmid into 293T cells for virus production. Cells were selected in 2.5 µg/mL puromycin in the culture medium or by fluorescence-activated cell sorting to generate the stable transfections.

### Cell proliferation assays

Cells were plated in 96-well plates and examined at 24, 48, 72, and 96 hours after plating (n=3). Cells were incubated with CellTiter 96 AQueous (MTS) solution for 3 hours. The absorbance at 490 nm was then measured using a microplate reader (BioTek).

### Cell scratch wound healing assay

Cells were plated at a density of 1×10^5^ cells/well in triplicate into 6-well plates. Once the cells had spread over the bottom of the wells, three or four parallel lines were scratched into each well using sterile 10μL tips. Suspended cells were washed off and remaining cells were cultured in medium without FBS. After 12 and 24 hours of incubation, for each well, five random fields were examined under a light microscope, photographed, and counted manually.

### Migration assay

Costar Trans-well migration plates with 8μm pore size (Corning, #3422) were pre-coated with Matrigel. Cells (1×10^5^) in 100μLRPMI medium without FBS were placed in triplicate into the upper chamber. To the lower chamber, 500 μL medium containing 10% FBS was added. After 12 and 24 hours of incubation, the plate inserts were removed and washed with PBS buffer several times to get rid of unattached cells. All the residual cells on the upper side were scraped with a cotton swab. Migrated cells on the lower side of the insert were fixed in 4% formalin for 20 minutes, washed twice with PBS, and stained with 0.1% crystal violet for 10 minutes. For each insert, five random fields were examined under a light microscope, photographed, and counted manually.

### *In vivo* xenograft assay

Cell suspensions (1×10^6^ cells) of cells, in a total volume of 100 μL mixed with Matrigel in a 1:1 ratio, were injected subcutaneously into the right flanks of 4-week-old male BALB/C nude mice (SLAC, Shanghai). The body weight and tumor volumes were measured and recorded every 3 days from 2 weeks after inoculation. Tumor volume was calculated with the following formula: volume= 0.5 × tumor length × tumor width^2^. Tumors were collected and photographed 60 days after inoculation. All mice were housed in the SPF animal facility in a pathogen-free environment with controlled temperature and humidity. All animal experiments were carried out following the ethical regulations of the Animal Care committee at Ren Ji Hospital.

### Statistical analysis

Statistical evaluation was conducted using Student’s t-test. Multiple comparisons were analyzed first by one-way analysis of variance. The log-rank (Mantel-Cox) test was used for patient survival analysis. The Pearson correlation was used to analyze the strength of the association between expression levels of Di-Ras2 and its related genes in patient samples. A significant difference was defined as P <0.05.

## Data availability statement

The data that support the findings of this study are available from the corresponding author upon reasonable request.

## Acknowledgements

This study was supported by funds from Ministry of Science and Technology of the People’s Republic of China (2017YFA0102900 to WQG), National Natural Science Foundation of China (81872406 and 81630073 to WQG, 81772938 to L. Li, 81602443 to X. Li), State Key Laboratory of Oncogenes and Related Genes (KF01801 to L. Li), Science and Technology Commission of Shanghai Municipality (16JC1405700 to WQG, 18140902700 and 19140905500 to L. Li), High Peak IV fund from Education Commission of Shanghai Municipality on Stem Cell Research (to WQG), KC Wong foundation (to WQG) and Hunan Provincial Natural Science Foundation of China (2019JJ50550 to X. Li). L. Li is supported by Innovation Research Plan from Shanghai Municipal Education Commission (ZXGF082101), and Shanghai Jiao Tong University Medical Engineering Cross Fund (YG2016MS52).

## Author contributions

HR mainly performed the experiments, analyzed the data and wrote the paper. XL collected the clinical samples. ML, JL, XL and JX helped with the experiments. LL and W-Q Gao carried out the experiment design and manuscript drafting. All authors had edit and approved the final manuscript.

---

**Figure S1.**
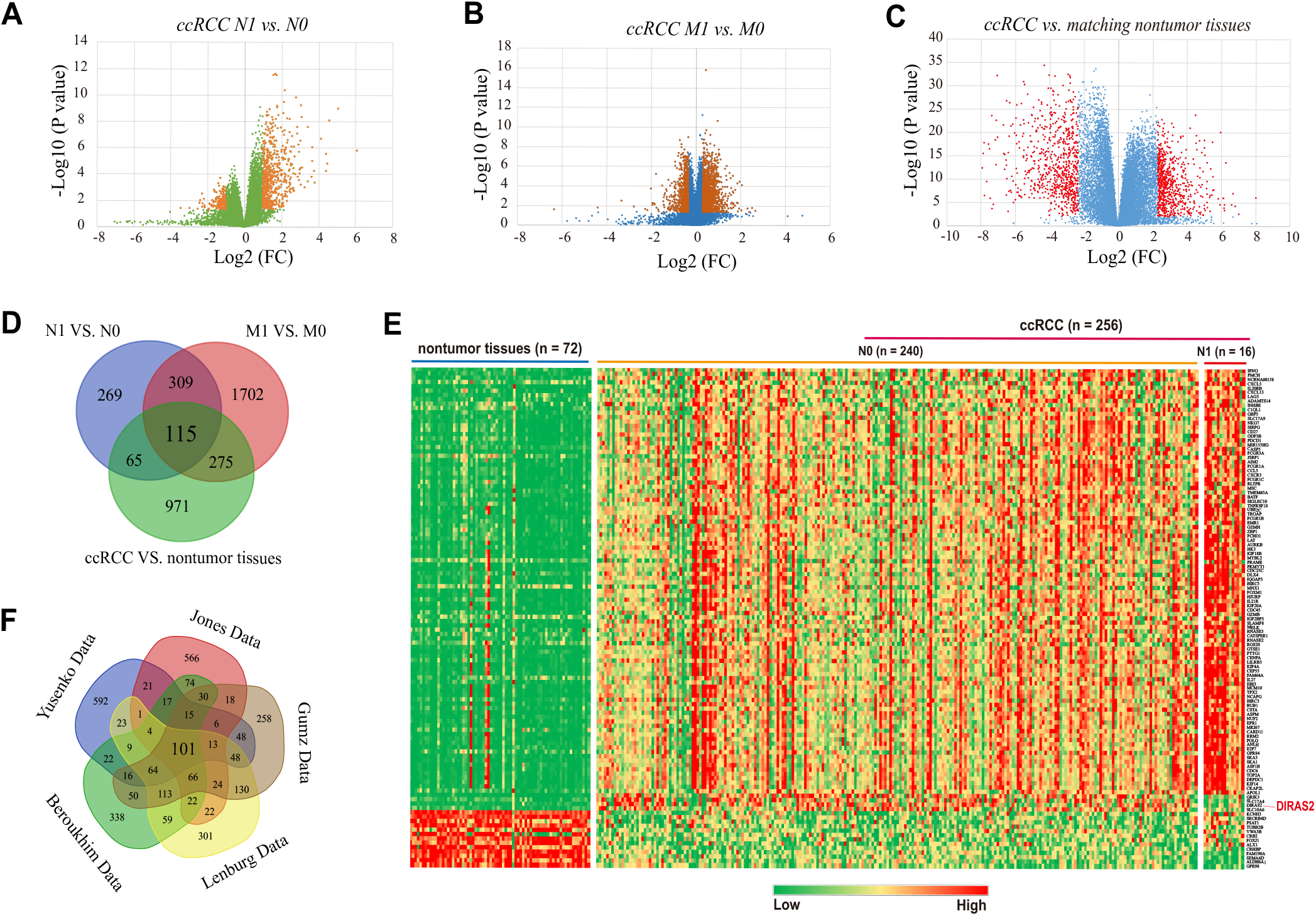
Differentially expressed genes in human kidney tumor samples. Venn diagram analysis of up-regulated genes based on the five independent datasets Yusenko Renal, Jones Renal, Gumz Renal, Beroukhim Renal, Lenburg Renal and from Oncomine database.

**Figure S2.**
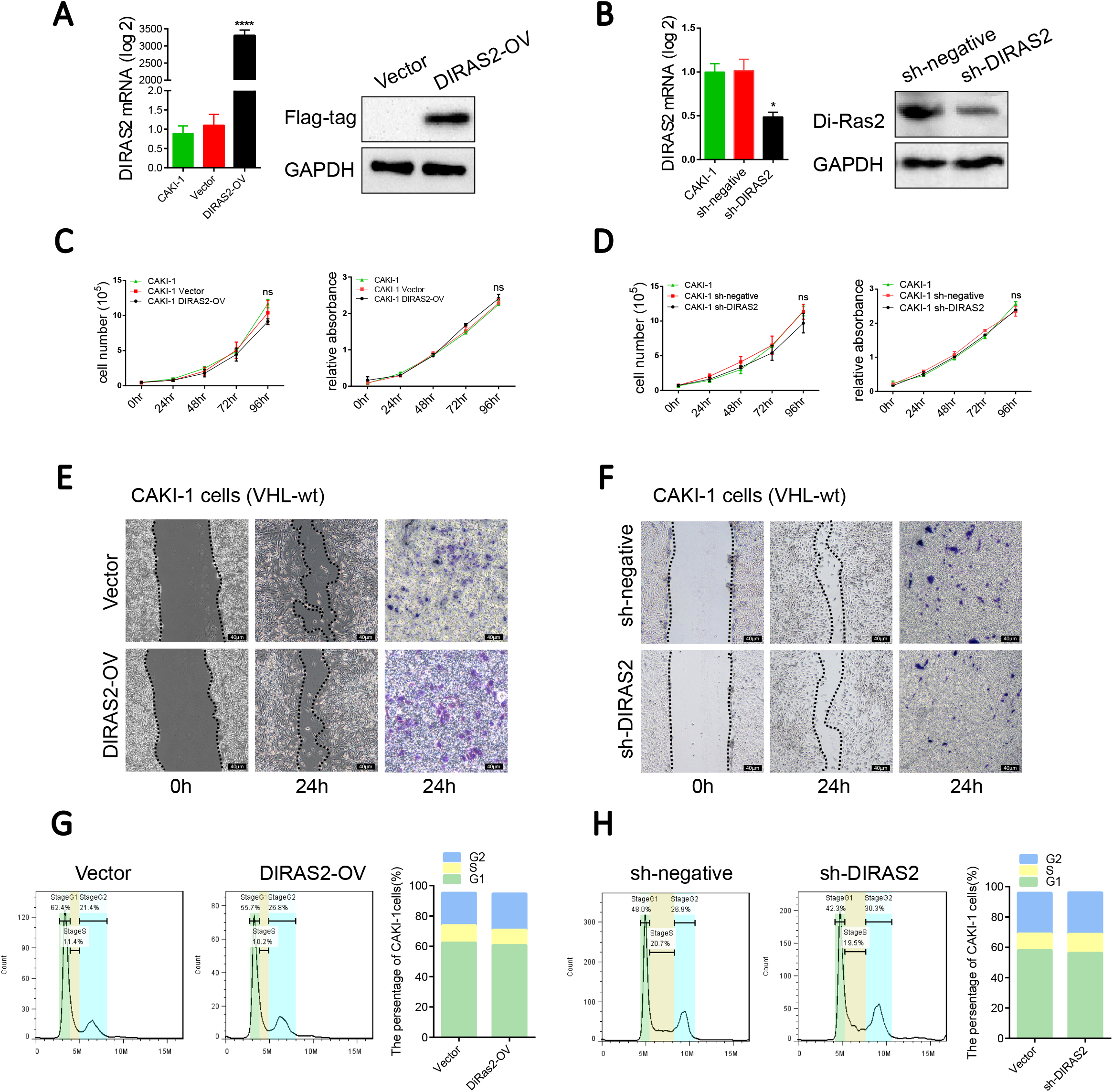
Di-Ras2 has no effect on VHL-wild-type CAKI-1 cells. mRNA and protein levels of Di-Ras2 after (**A**) overexpression and (**B**) knockdown of Di-Ras2 in CAKI-1 cells lines. (**C, D**) Cell proliferation abilities were compared with their vector control cells. Cell migration and invasion abilities of (**E**) DIRAS2-OV CAKI-1 cells as well as (**F**) DIRAS2-KD CAKI-1 cells were compared with their vector control cells, as demonstrated by cell wound scratch and trans-well assays. The percentages of (**G**) DIRAS2-OV CAKI-1 cells as well as (**H**) DIRAS2-KD CAKI-1 cells in the G1, S, and G2 phases for each sample are shown. VT, vector; WT, wild-type; MUT, mutated; OV, overexpression; KD, knockdown. Data are shown as the mean value ± SEM.

**Figure S3.**
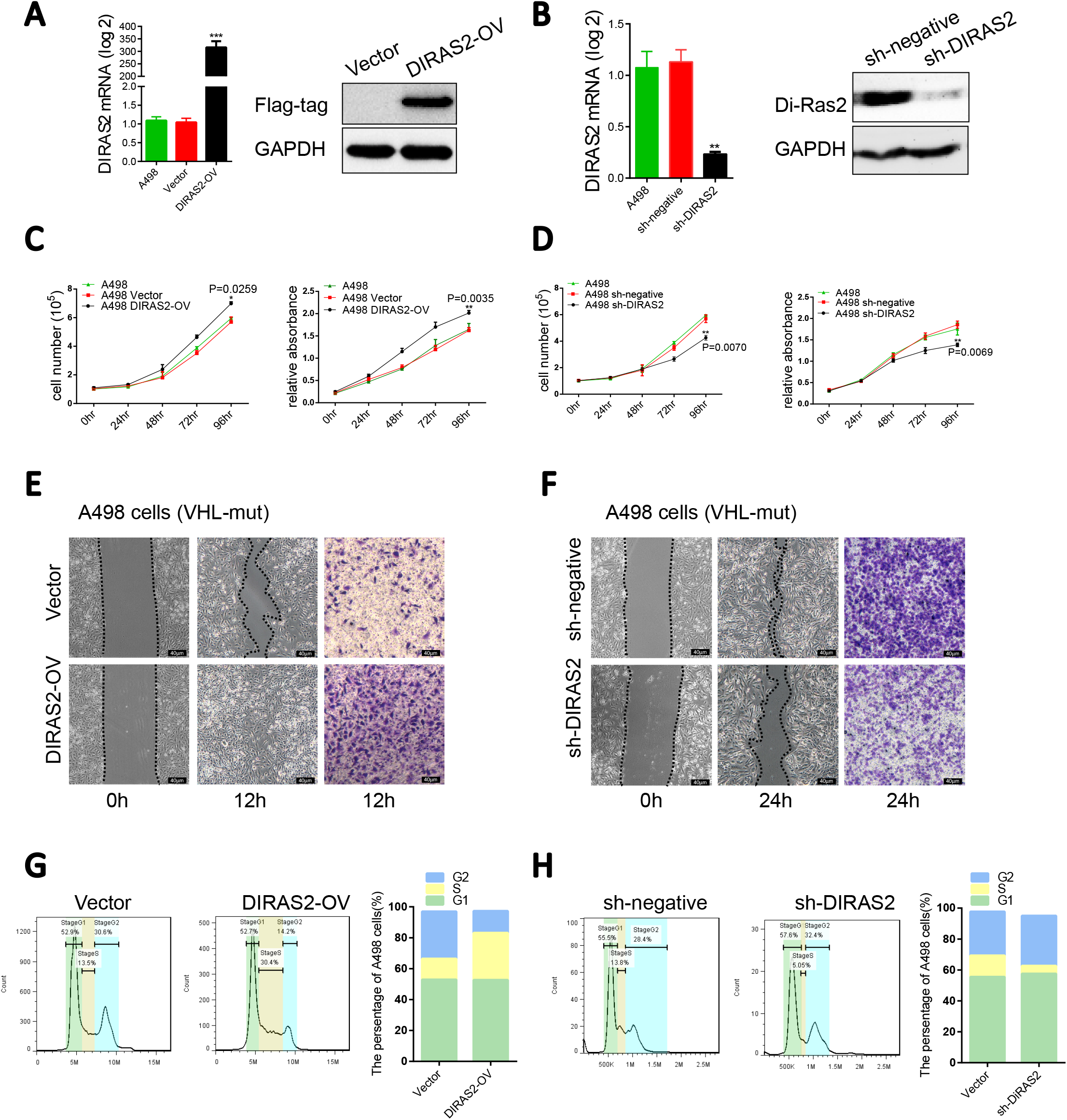
Di-Ras2 promotes ccRCC cell proliferation, migration, and invasion in VHL-mutated A498 cells. mRNA and protein levels of Di-Ras2 after (**A**) overexpression and (**B**) knockdown of Di-Ras2 in A498 cells lines. (**C, D**) Cell proliferation abilities were compared with their vector control cells. Cell migration and invasion abilities of (**E**) DIRAS2-OV A498 cells as well as (**F**) DIRAS2-KD A498 cells were compared with their vector control cells, as demonstrated by cell wound scratch and trans-well assays. The percentages of (**G**) DIRAS2-OV A498 cells as well as (**H**) DIRAS2-KD A498 cells in the G1, S, and G2 phases for each sample are shown. VT, vector; WT, wild-type; MUT, mutated; OV, overexpression; KD, knockdown. Data are shown as the mean value ± SEM.

**Figure S4.**
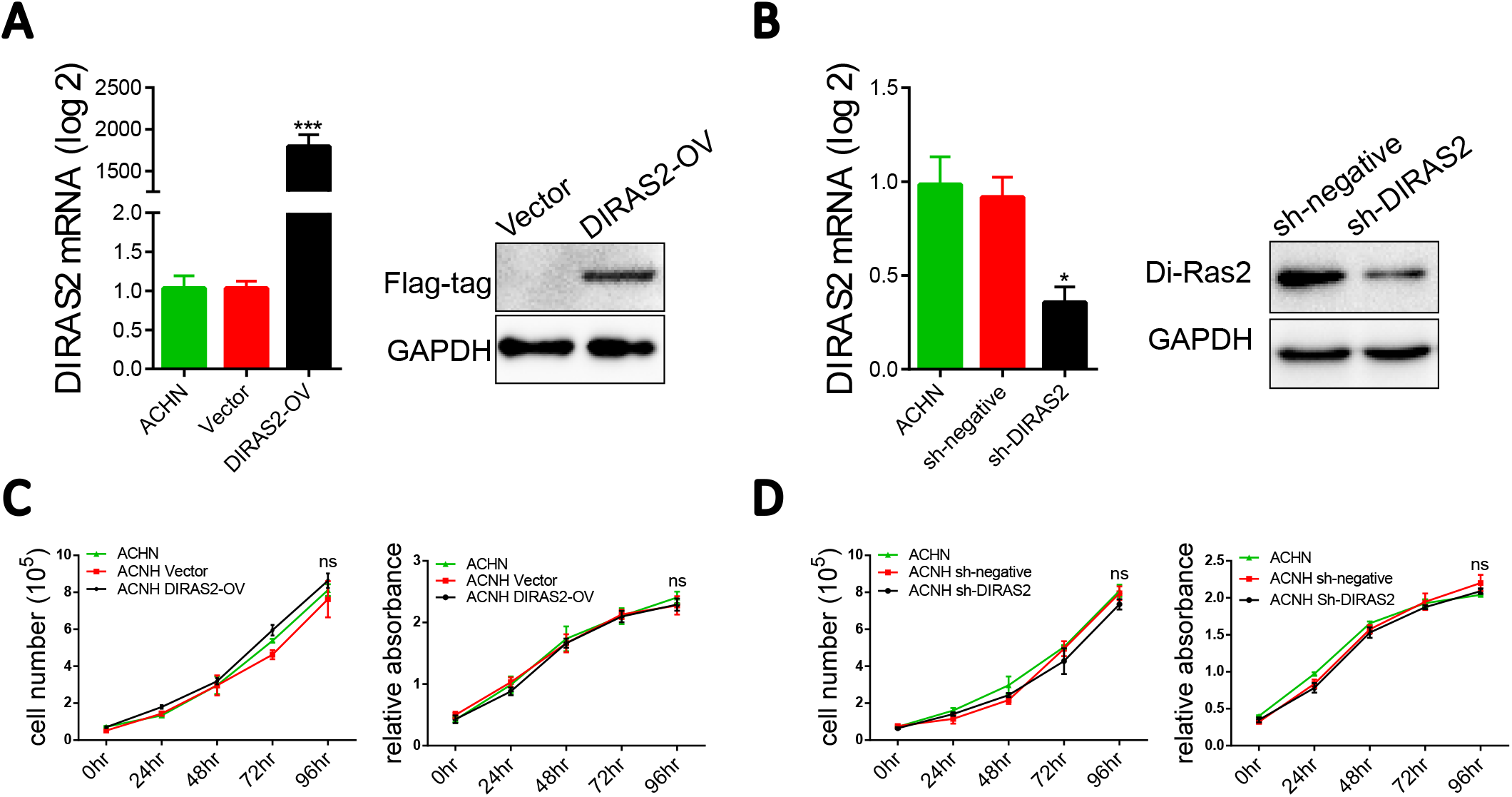
Di-Ras2 has no effect on VHL-wild-type ACHN cells. mRNA and protein levels of Di-Ras2 after overexpression (**A**) and knockdown (**B**) of Di-Ras2 in ACHN cells. (**C, D**) Cell proliferation abilities were compared with their vector control cells. VT, vector; WT, wild-type cells; MUT, VHL-mut cell; OV, overexpression Di-Ras2 lentivirus-infected cells; KD, shRNA lentivirus-infected cells. Data are shown as the mean value ± SEM.

**Figure S5.**
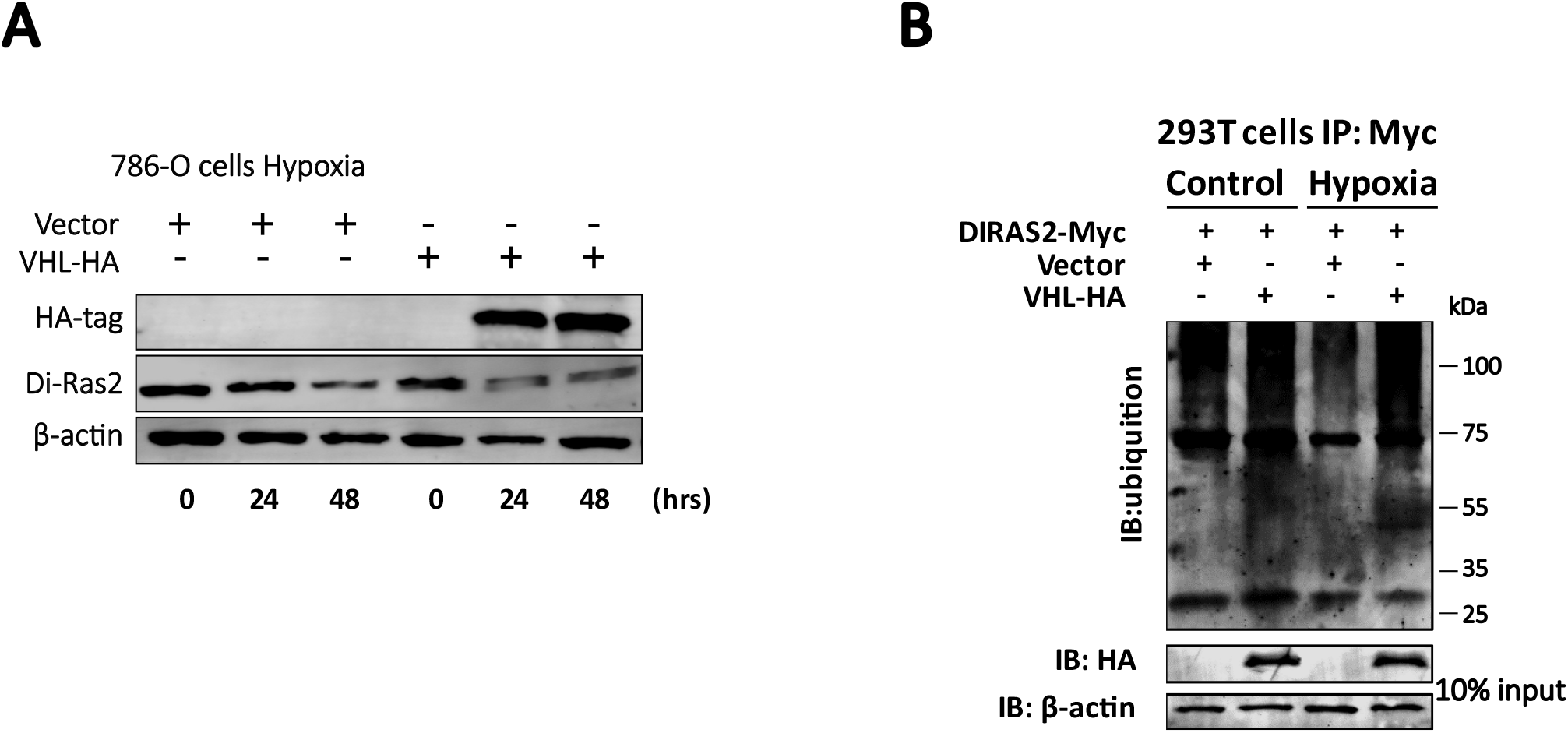
VHL mediated ubiquitination of Di-Ras2 is independent of hypoxia. (**A**) The protein abundance of Di-Ras2 in 786-O cells under hypoxia culture. (**B**) Ubiquitination of exogenous Di-Ras2 were enhanced by VHL overexpression in 293T cells and CAKI-1 cells under both normal and hypoxia culture.

